# Selfish grower behavior can group-optimally eradicate plant diseases caused by coinfection

**DOI:** 10.1101/2023.11.19.567700

**Authors:** Frank M. Hilker, Lea-Deborah Kolb, Frédéric M. Hamelin

**Affiliations:** Institute of Mathematics and Institute of Environmental Systems Research, Osnabrück University, Barbarastr. 12, D–49076 Osnabrück, Germany; Department of Ecosystem Services, Helmholtz Centre for Environmental Research–UFZ, Institute of Biodiversity, Friedrich Schiller University Jena, and German Centre for Integrative Biodiversity Research (iDiv) Halle–Jena–Leipzig, Puschstr. 4, D–04103 Leipzig, Germany; Institut Agro, Univ. Rennes, INRAE, IGEPP, 35000, Rennes, France

**Keywords:** Infectious disease, human behavior, mathematical model, epidemiological game, imitation dynamics

## Abstract

Game-theoretic behavioral and epidemiological models suggest that it is impossible to eradicate a disease through voluntary control actions when individuals behave according to their own interests. The simple reason is that free-riding behavior, which is best for self-interest, leads to a control coverage on the group level that is insufficient to eradicate the disease. Here we show that, for diseases that are caused by coinfection, self-interest can actually result in the socially optimal outcome of disease eradication. Our result challenges the conventional wisdom that selfish behavior undermines the group interest; it resolves a social dilemma in the absence of any cooperation, institutional arrangements, or social norms. Our model is motivated by coinfecting plant viruses, common among agricultural crops and natural plants, and the behavioral dynamics of growers to adopt protective action (biological or chemical control). The epidemiological scenario, in which the disease is eradicated by self-interest, is characterized by a positive feedback process in which coinfection enhances infectivity. Similar feedback structures exist for a range of typical epidemiological processes that facilitate disease persistence if prevalence is sufficiently large. The underlying mechanism may therefore be applicable to other diseases.

## 1. Introduction

Human behavior plays an important role in the control of infectious diseases (Funk et al., 2010; Schaller, 2011; Bauch & Galvani, 2013; Verelst et al., 2016; Chang et al., 2020; Bedson et al., 2021). Individual decision-making, e.g., on whether or not getting vaccinated, adhering to social distancing, or wearing face masks, can significantly affect disease spread on a population level. It is a basic tenet of epidemiology that the successful control of disease spread requires a sufficiently large fraction of the population adopting disease control (Anderson & May, 1982; Fine, 1993; Keeling & Rohani, 2007; Stone et al., 2012). This is probably most famously expressed by the “herd immunity” threshold, which translates into disease eradication as long as a critical proportion of the population has been afforded protection, e.g., in the form of vaccination.

In fact, it is not required that all individuals engage in disease control, because they can be indirectly protected by the action of others. By adopting protective measures, an individual reduces the risk of infection, but this comes with some financial expense, possibly social cost, or even health risk. Alternatively, by avoiding protective measures, an individual can spare the costs but still benefit from avoiding infection when the other individuals and potential contacts have engaged in protective measures. The latter behavior is also known as “free-riding” and impedes the efforts toward disease control. Voluntary control measures can be understood as a social dilemma (or collective action problem, Siegal et al., 2009), as the self-interest of an individual may not be the best option for the population (Fine & Clarkson, 1986). The self-limiting effect of the participation in protective measures may be a major reason for the failure of some disease eradication programs (Geoffard & Philipson, 1997). Indeed, the recent review by Chang et al. (2020) has revealed that, according to all models accounting for human behavior, individual self-interest prevents disease eradication, unless some specific conditions are met.

Here, we consider diseases that are caused by coinfection of hosts by two or more pathogen species or strains of the same species (Balmer & Tanner, 2011; Vaumourin et al., 2015; Hamelin et al., 2019). Coinfection by multiple viruses is widespread in agricultural crops (Allen et al., 2019; Moreno & López-Moya, 2020) and natural plant communities (Seabloom et al., 2009; Susi et al., 2015). Often, infection by only one virus results in rather mild symptoms, but coinfection can be particularly damaging. Plant diseases caused by coinfection are a main threat to global food security and human health (Reynolds et al., 2015; Rybicki, 2015). Examples include maize lethal necrosis (caused by *Maize chlorotic mottle virus* and a virus from the family *Potyviridae*; Redinbaugh & Stewart, 2018), sweet potato virus disease (caused by *Sweet potato feathery mottle virus* and *Sweet potato chlorotic stunt virus*; Kokkinos et al., 2006), and rice tungro disease (caused by *Rice tungro bacilliform virus* and *Rice tungro spherical virus* coinfection; Hibino et al., 1978).

A notorious feature of coinfection is that the presence of two viruses can enhance the infectivity and thus promote transmission of the viruses (Alcaide et al., 2020; Lamichhane & Venturi, 2015). This induces a self-reinforcing feedback loop: The more coinfections, the higher the transmission rates. If neither virus by itself is able to invade the host population, the self-reinforcing feedback can nevertheless facilitate the spread of the viruses, provided the prevalence of coinfection is sufficiently large (Allen et al., 2019; Keeling & Rohani, 2007, Sect. 4.1.3). This creates a bistable situation, mathematically related to a backward bifurcation. The eradication of the disease is particularly difficult in this case, because it is not sufficient when control measures reduce the basic reproduction number below one. Instead, control must be more intensive to reduce the basic reproduction number even further because of the coinfection-induced bistability.

The management of plant diseases and the understanding of control successes or failures increasingly relies on dynamic modeling (Cunniffe et al., 2016; Hamelin et al., 2021; Ristaino et al., 2021). The extension of epidemiological models beyond single viruses is considered a key challenge in modeling plant diseases (Cunniffe et al., 2015, challenge 4). Moreover, while there is an increasing number of epidemiological models accounting for human behavior and socio-economic feedbacks in the control of human (e.g., Bauch et al., 2003; Galvani et al., 2007; Fenichel et al., 2011; Cascante-Vega et al., 2022) and animal diseases (e.g., Hidano et al., 2018; Delabouglise & Boni, 2020; Cristancho Fajardo et al., 2021), there are only a few studies for plant diseases (Milne et al., 2015, 2020; McQuaid et al., 2017; Bate et al., 2021; Saikai et al., 2021; Murray-Watson et al., 2022; Murray-Watson & Cunniffe, 2022, 2023). However, their model structures are relatively complex and obscure how the epidemiological and behavioral processes interrelate (see Jeger et al., 2023).

In this paper, we first develop a very simple epidemiological model that captures the essence of coinfection-induced infectivity enhancement and that results in bistability between a disease-free and an endemic coinfection equilibrium. We then extend this coinfection model to include dynamic grower behavior regarding the use of pesticides or natural enemies (i.e., chemical or biological control) as a protective measure of disease control. Grower behavior is described by imitation dynamics (Hofbauer & Sigmund, 1998; Helbing, 2010), a social learning process in which individual growers adopt the action of growers with better pay-off. We show that, in the epidemiological regime of infectivity-enhanced bistability, self-interest can actually lead to successful disease eradication.

## 2. Model description

We begin with describing the epidemiological dynamics of the model. Later, we will add the behavioral dynamics and arrive at the integrated behavioral-epidemiological model.

### 2.1 Coinfection model

The derivation of the coinfection model follows the approach in Hamelin et al. (2019). We consider a compartment model for a host plant population and two virus strains. Let *J*_∅_ ≥ 0 be the density of host plants that carry neither virus, *J*_1_ ≥ 0 and *J*_2_ ≥ 0 the densities of host plants that carry exclusively virus 1 and virus 2, respectively, and *J*_1,2_ the density of host plants that are coinfected by both viruses and thus develop the disease. The epidemiological dynamics are given by

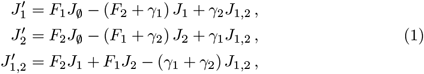

where the prime denotes differentiation with respect to time. *γ*_1_, *γ*_2_ *>* 0 are the per-capita recovery rates from infection with virus 1 and 2, respectively. The total host population density *N* = *J*_∅_ + *J*_1_ + *J*_2_ + *J*_1,2_ remains constant. *F*_1_ and *F*_2_ are the forces of infection of virus 1 and 2, respectively:

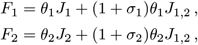

where *θ*_1_, *θ*_2_ *>* 0 are the horizontal transmission coefficients of virus 1 and 2, respectively. Coinfected hosts are assumed to have a modified infectivity compared to hosts that carry exclusively one virus. This is described by the relative infectivities *σ*_1_, *σ*_2_ *>* −1. If *σ*_1_ *>* 0, then coinfected hosts have an enhanced infectivity of virus 1 compared to hosts infected exclusively by virus 1. If −1 *< σ*_1_ *<* 0, then coinfected hosts have a diminished infectivity of virus 1 compared to hosts infected exclusively by virus 1. If *σ*_1_ = 0, then there is no effect of coinfection on infectivity of virus 1. Analogously for *σ*_2_.

Let *I*_1_ = *J*_1_ + *J*_1,2_ and *I*_2_ = *J*_2_ + *J*_1,2_ be the densities of host plants that have been infected (not necessarily exclusively) by virus 1 and virus 2, respectively. Then the forces of infection can be written as

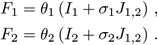

*N* − *I*_1_ and *N* − *I*_2_ are the host densities susceptible to infection by virus 1 and 2, respectively. System (1) can then be written as

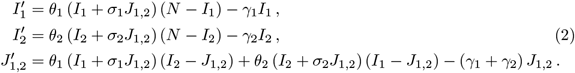

We now show that the coinfected density *J*_1,2_ can be substituted by the product *I*_1_*I*_2_*/N* . Let *Z* = *I*_1_*I*_2_*/N* − *J*_1,2_. Then

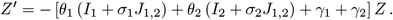

Since the expression in the brackets is strictly positive, *Z*(*t*) → 0 as *t* → ∞. Consequently, *J*_1,2_ → *I*_1_*I*_2_*/N* as *t* → ∞. Therefore, system (2) asymptotically reduces to a two-dimensional system. Replacing *J*_1,2_ by *I*_1_*I*_2_*/N*, we obtain

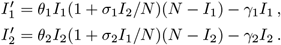

For simplicity, let us assume that the epidemiological parameters do not differ between the two viruses, i.e.,

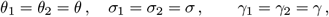

which yields the symmetric model

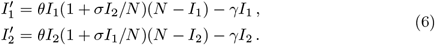

### 2.2. Behavioral model

Let us now introduce disease management and the behavioral dynamics of growers. Control options taken will depend on the transmission pathways of the disease. In general, there are various routes of transmission. For example, the viruses causing maize lethal necrosis are transmitted both horizontally by insects and vertically via seed, and there is also evidence for soil-borne transmission (see Hilker et al., 2017, and references therein). There is little information on the management practises most often applied to control diseases caused by coinfection. However, many plant viruses are vectored by insects such as aphids, thrips, and whiteflies (Eigenbrode et al., 2018). Pesticides reduce vector populations and, thus, virus transmission. Chemical (or biological) control therefore emerges as a control option for growers with the sufficient financial assets (Strange & Scott, 2005) and will be the one considered here.

We assume that growers always choose between two strategies *X* and *¬X*, namely to apply pesticides as a form of disease control or not to apply them, respectively. Along the game dynamic approach by Bauch (2005), growers adopting disease control are assumed to receive the payoff

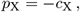

where *c*_X_ *>* 0 are the costs of purchasing and applying pesticides. The perceived payoff for growers who do not adopt disease control depends on the cost *c*_I_ *>* 0 of diseased plants (e.g., lost harvest or lower crop quality) and on the perceived risk of disease:

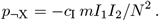

The perceived risk of disease is proportional to the disease prevalence (*I*_1_*I*_2_*/N* ^2^), where a higher value of *m >* 0 means that growers perceive the disease as more harmful. Note that we assume that disease damage only occurs due to coinfection and that infection with a single virus causes no notable reduction in payoff.

In the proportional imitation dynamics (Cressman, 2003), growers are assumed to sample each other randomly at rate *ψ >* 0. If the payoff of the sampled grower is higher than one’s own payoff, the strategy of the sampled grower is adopted with a probability proportional to the expected gain in payoff. The payoff gain from switching to disease control is

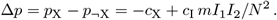

The fraction of controllers in the grower population changes according to the following equation well-known from imitation dynamics (e.g., Bauch, 2005):

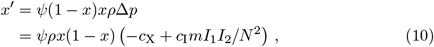

where *ρ* is the constant of proportionality between switching probability and expected payoff gain.

### 2.3 Coupled model

Coupling the coinfection model (6) with the behavioral model (10), we obtain

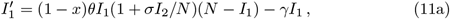

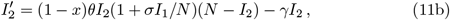

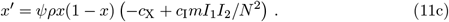

In (11a) and (11b), we multiply the terms accounting for new infections with (1 − *x*). This represents the effect of growers using pesticides on the epidemiological dynamics. There are a number of implicit assumptions in this model formulation. First, pesticides affect the transmission pathways of the two viruses equally. This certainly holds for coinfecting viruses that share the same vector species, of which there are many examples (e.g., Lacroix et al., 2014; Salvaudon et al., 2013; Holt & Chancellor, 1996). Although there are also many examples where coinfecting viruses have different vector species (e.g., Peñaflor et al., 2016), the pesticide may still be assumed equally effective in these cases. Second, growers who use pesticides in their fields are assumed to sufficiently reduce the local vector population to prevent new infections. Assuming growers own fields of equal area, the reduction in the overall increase in infection is proportional to the fraction of growers employing pesticides.

Let us now nondimensionalize system (11). We introduce

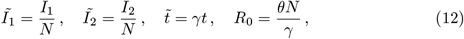

and

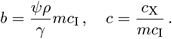

Parameter *R*_0_ is the basic reproduction number (which is identical for the two viruses due to the assumption of virus symmetry). Parameters *b* and *c* are composite socio-economic parameters. They both include the term *mc*_I_. In the remainder, we shall refer to *c* as *relative costs* because it is additionally influenced by the costs of disease control, *c*_X_. We shall refer to *b* simply as *scaled learning rate* because it is additionally influenced by the imitation parameters *ψρ*.

Dropping the tildes in (12) for notational convenience, system (11) becomes

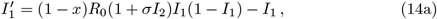

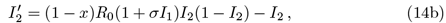

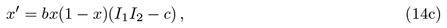

where the prime now denotes differentiation with respect to time, which has been scaled relative to the period of virus infectiousness. This is the model that we will analyze in this paper. We shall henceforth refer to it as coupled model, but we note that models of this type are also called “behavior-prevalence” models (e.g., Bauch et al., 2013). All parameters are strictly positive, except *σ >* −1. Note that all three state variables are fractions, i.e., the proportion of host plants infected with virus 1 or 2, or the proportion of growers applying disease control. That is, the state space of the model is (*I*_1_, *I*_2_, *x*) ∈ [0, 1]^3^. The structure of the model is illustrated in Fig. 1. The formulation of the model reveals that the fraction of controllers among the growers increases if and only if the perceived disease risk (represented by the prevalence *I*_1_*I*_2_) outweighs the relative costs of disease control (*c*), see Eq. (14c).

**Fig. 1.**
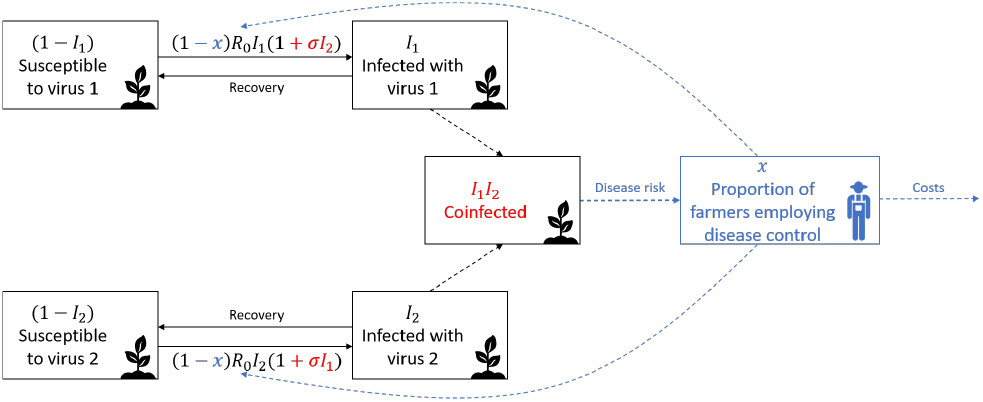
Illustration of the coupled behavioral-epidemiological model (14). Dashed arrows indicate that the disease prevalence is given by the product of coinfected plants, which influences the proportion of farmers applying control, which in turn feeds back onto the virus transmission.

## 3 Results on the coinfection model

### 3.1 No control

In the absence of control, i.e., *x* = 0, system (14) simplifies to

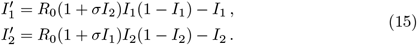

This system has been analyzed in Appendix A. The dynamics depend only on two epidemiological parameters and can be summarized as follows (see Fig. 2). If *R*_0_ *>* 1, both viruses spread in the population and establish a stable unique endemic equilibrium in which both viruses coexist. If *R*_0_ *<* 1, the disease-free equilibrium is locally asymptotically stable. However, if and only if *σ >* 1 and

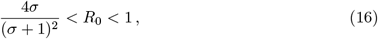

then there is a bistable scenario in which both the disease-free equilibrium and the larger of two endemic equilibria are locally stable. That is, the disease may persist even though *R*_0_ *<* 1, provided the infectivity enhancement, i.e. *σ*, is sufficiently large.

**Fig. 2.**
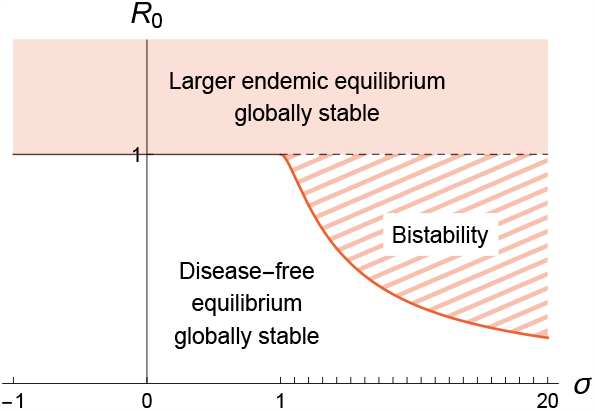
Dynamical regimes of the coinfection model (15) without control. In the bistable region, the two attractors are the disease-free equilibrium and the larger endemic equilibrium. The red curve corresponds to saddle–node bifurcations and is given by condition (16). The dashed (solid) black curves are backward (transcritical, respectively) bifurcations.

### 3.2 Constant control

If *x* is not a dynamic state variable but a constant parameter, system (14) simplifies to

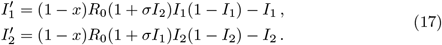

Following basic epidemiological theory (e.g., Keeling & Rohani, 2007), the minimum control coverage to eradicate the disease (i.e., for the disease-free equilibrium to be globally asymptotically stable) is

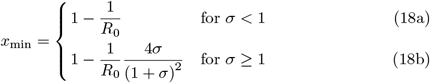

This is the constant control coverage needed for “herd immunity”. The expression in (18a) is the same as the one required for voluntary vaccination (e.g., Keeling & Rohani, 2007); see the yellow surface in Fig. 3. The expression in (18b) is the one required because of coinfection effects; see the red surface in Fig. 3. Note that the coinfection effects elevate the minimum control coverage.

**Fig. 3.**
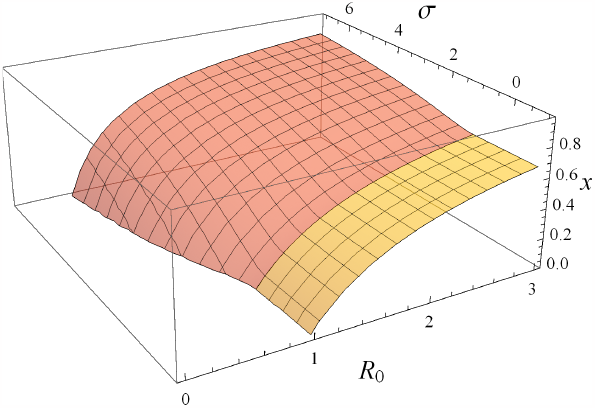
Minimum control coverage required for disease elimination in the model with constant control (17). The yellow and red surfaces are given by (18a) and (18b), respectively.

## 4. Results on the coupled model

We now turn to the coupled coinfection-behavior model in which the fraction of controllers among the growers changes dynamically. Table 1 summarizes the results of the mathematical analysis. The details are given in Appendix B. An important role is played by 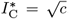; it represents the minimum infection levels of *I*_1_ and *I*_2_ required for disease control to become worthwhile for the growers. That is, the control coverage *x* increases for 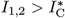 (or *I*_1_*I*_2_ *> c*) and decreases for 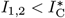 (or *I*_1_*I*_2_ *< c*).

**Table 1.** Equilibria 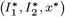 of system (14) and their existence and local stability conditions.

| Description | Components | Existence | Stability |
| --- | --- | --- | --- |
| Disease-free | $E_0 = (0, 0, 0)$ | always | $R_0 < 1$ |
| Full control | $E_F = (0, 0, 1)$ | always | never |
| Mono-pathogenic virus 1 | $E_1 = \left(1 - \frac{1}{R_0}, 0, 0\right)$ | $R_0 > 1$ | never |
| Mono-pathogenic virus 2 | $E_2 = \left(0, 1 - \frac{1}{R_0}, 0\right)$ | $R_0 > 1$ | never |
| Endemic (small) | $E_- = (I_-^*, I_-^*, 0)$ | Tab. 2 | never |
| Endemic (large) | $E_+ = (I_+^*, I_+^*, 0)$ | Tab. 2 | $I_+^* < \sqrt{c} = I_C^*$ |
| Control | $E_C = (\sqrt{c}, \sqrt{c}, x_C^*)$ | $\sigma \geq \sigma_C$ | $\sigma < \sigma_{HB} = \frac{1}{1-2\sqrt{c}}$ |

*Note:* $I_-^*$ and $I_+^*$ are defined in Eq. (A.3). $x_C^*$ is defined in Eq. (B.11). $\sigma_C$ is defined in Eq. (B.12).
note that it depends only on the relative costs $c$ . We can distinguish the following dynamical regimes.

**Table 2.** Summary of the existence conditions of the smaller and larger endemic equilibria as a function of parameters *σ* and *R*_0_. The conditions apply equally to the endemic equilibria 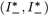 and 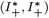 in the coinfection model, and to the endemic equilibria *E*_−_ and *E*_+_ in the coupled model.

| | $-1 < \sigma < 0$ | $0 < \sigma < 1$ | $\sigma > 1$ |
| --- | --- | --- | --- |
| $R_0 > 1$ | smaller | larger | |
| $R_0 < 1$ | none | | smaller and larger, iff $R_0 > \frac{4\sigma}{(\sigma+1)^2}$ |

### 4.1 Bistable epidemiological scenario

The phase portraits in Fig. 4 illustrate the dynamics of the coupled model, when the coinfection model is bistable (i.e., *R*_0_ *<* 1). 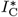 is marked by a blue vertical line; note that it depends only on the relative costs *c*. We can distinguish the following dynamical regimes.

A. If control is too expensive (large value of *c*, Fig. 4A), the blue vertical line is located to the right of the larger endemic equilibrium. That is, the disease risk perceived by the growers does not outweigh the high costs of control, and nobody engages in control (*x*^∗^ = 0). The coupled system still exhibits bistability between the disease-free equilibrium (DFE) and the larger endemic equilibrium. The separatrix between the basins of attraction emanates from the smaller endemic equilibrium; with increasing control coverage the DFE expands its basin of attraction compared to the larger endemic equilibrium.
B. If the relative costs are reduced such that 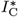 is between 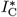 and the hump of the orange curve (Fig. 4B), a stable control equilibrium establishes itself. That is, disease control is now sufficiently affordable for the given disease risk. The system remains bistable, but one of the attractors is now the control equilibrium which has lower infection levels than the larger endemic equilibrium. Moreover, the basin of attraction to the DFE is further expanded compared to that in Fig. 4A.
C. If 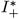 is between the the hump of the orange curve and 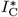 (Fig. 4C), the control equilibrium is unstable and surrounded by a limit cycle. That is, low costs can destabilize the control equilibrium. If the costs of disease control become more affordable, more growers adopt control such that the disease prevalence decreases. In turn, this disincentivizes growers from employing control with the effect of disease prevalence going up again, and the cycle starts anew. The system is still bistable, with the other attractor being the DFE. Note that the amplitudes of the limit cycle increase with lower relative costs. That is, the limit cycle approaches more and more the basin of attraction to the DFE as *c* decreases (Fig. 4D).
D. When the relative costs are further reduced such that 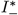 moves further to the left (Fig. 4D), the limit cycle has disappeared in a homoclinic bifurcation (as it collides with the smaller endemic equilibrium). Now, the DFE is the only attractor and globally asymptotically stable. That is, long-term disease eradication is guaranteed. However, there may be transient oscillations for initial conditions in the vicinity of the unstable control equilibrium; they trace the remnants of the disappeared limit cycle but eventually vanish toward the DFE.
E. If 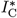 is below 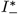 (Fig. 4E), there is no control equilibrium. This is because the control has reduced the infection levels into the basin of attraction of the DFE. The DFE is, therefore, the only attractor such that there is again successful disease eradication.

**Fig. 4.**
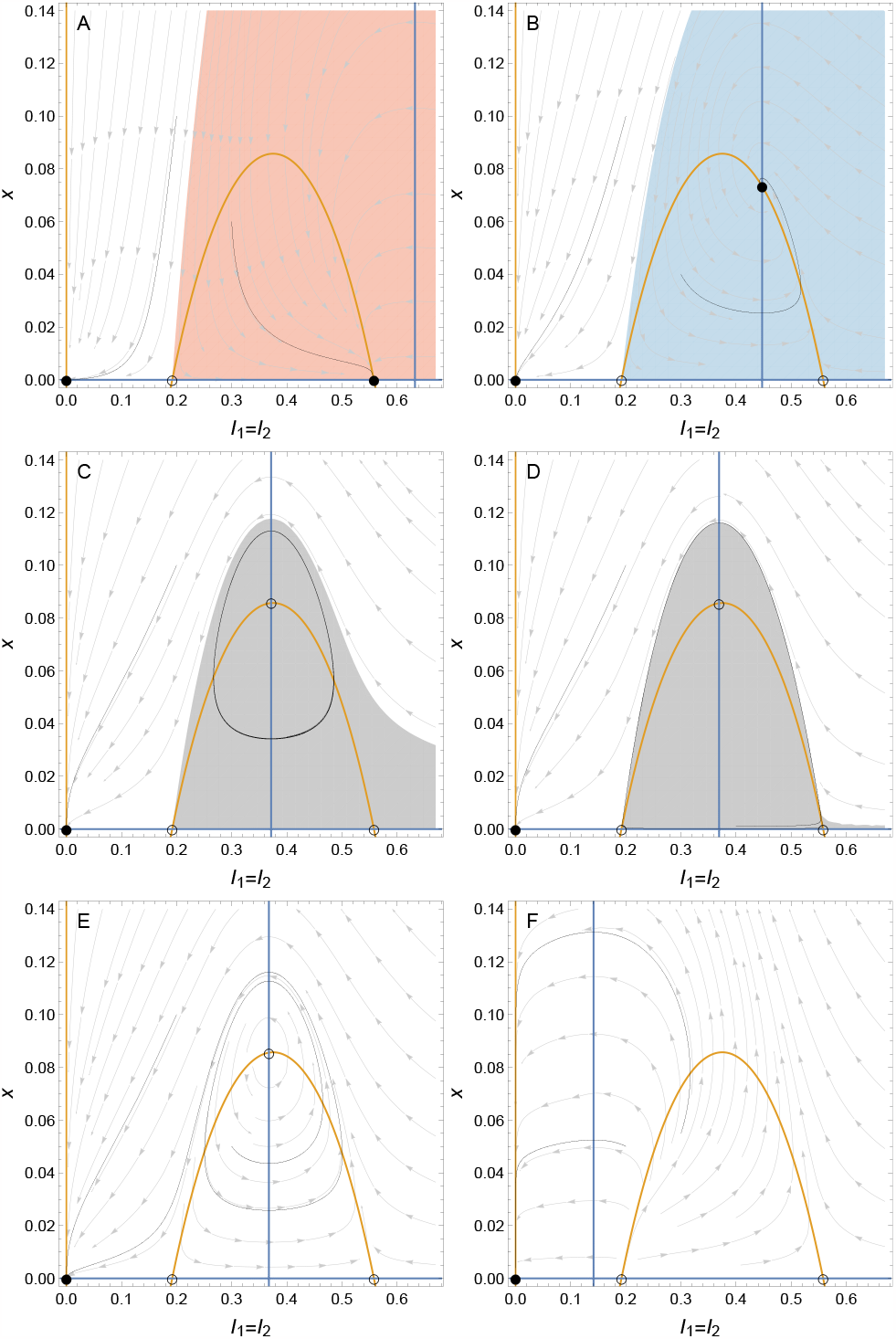
Phase portraits of the coupled model (14) for different relative control costs *c*. The threedimensional phase space is projected into the phase plane (*I, x*), where the infection levels *I*_1_ = *I*_2_ = *I* are set equal because of identical initial infection levels *I*_1_(0) = *I*_2_(0). Blue lines mark the zero-growth curves of *x* and orange lines mark the zero-growth curves of *I*. Full (empty) symbols mark stable (unstable) equilibria. Black thin lines are sample trajectories. The white areas are the basins of attraction to the disease-free equilibrium, red to the larger endemic equilibrium, blue to the control equilibrium, and gray to the limit cycle. Parameter values: *R*_0_ = 0.7, *σ* = 4, *b* = 1, (A) *c* = 0.4, (B) *c* = 0.2, (C) *c* = 0.138, (D) *c* = 0.1362452, (E) *c* = 0.135, (F) *c* = 0.02.

The bifurcation diagrams in Fig. 5 show how the long-term control uptake and infection levels change as a function of the relative costs. If the costs are too high (approx. *c >* 0.31), no control will be adopted. At *c* ≈ 0.31, there is a transcritical bifurcation, in which the control equilibrium establishes itself and exchanges stability with the larger endemic equilibrium. Inexpensive control destabilizes the system: at *c* ≈ 0.14, there is a Hopf bifurcation, which renders the control equilibrium unstable. This takes place at the local maximum of the control uptake, for details see Appendix B.3. The limit cycle that emerges in the Hopf bifurcation exist only in a small range of parameter values for *c*. The limit cycle disappears in a homoclinic bifurcation, which occurs when the amplitudes of the limit cycle “touch” either of the endemic equilibria. For *c <* 0.136, approximately, the coupled system is diseasefree as the DFE is the only attractor. Note that the control equilibrium disappears in another transcritical bifurcation at *c* ≈ 0.037, but this does not change that the DFE is globally asymptotically stable.

**Fig. 5.**
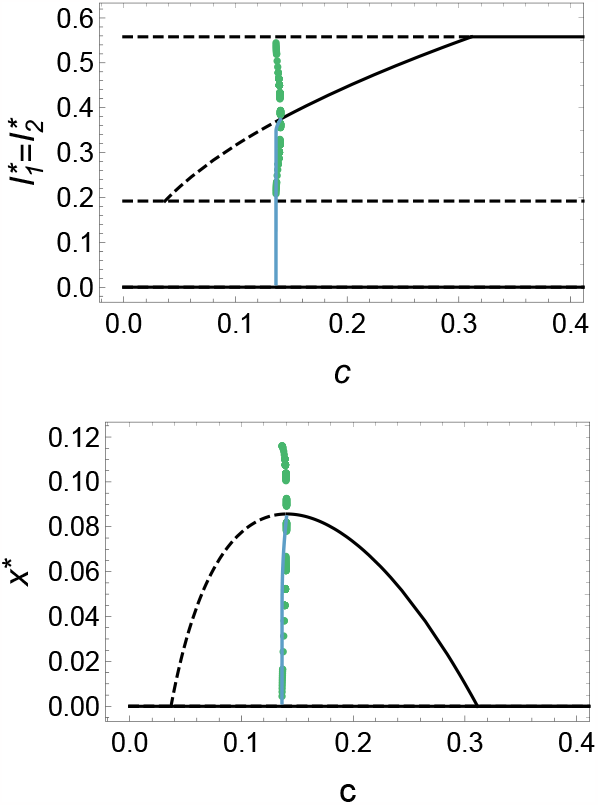
Bifurcation diagrams of the coupled model (14), when the coinfection system is bistable in the absence of control. Solid (dashed) lines represent stable (unstable) equilibria. Green points are the maxima and minima of stable limit cycles. The blue curves give the time averages along limit cycles. Parameter values: *R*_0_ = 0.7, *σ* = 4, *b* = 1.

Overall, we observe that disease eradication will be achieved for *c <* 0.136, approximately. For more expensive controls, the disease may persist depending on the initial conditions as the coupled system is bistable in various forms. One of the attractors is always the DFE; the alternative attractors are either the larger endemic equilibrium, the control equilibrium, or the limit cycle. Note that the infection levels 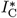 at the control equilibrium follow a square root branch and decrease with decreasing *c*. This trend also holds for the time-averaged infection levels during limit cycle oscillations. The control uptake at equilibrium initially increases as control becomes more affordable. However, once the Hopf bifurcation has taken place, the time-averaged control uptake decreases with decreasing *c*.

Figure 6A shows a two-parameter bifurcation diagram when additionally the infectivity enhancement *σ* is being varied. If *σ* is too small (approx. *σ <* 3.4), infectivity enhancement is not sufficient to facilitate disease spread because *R*_0_ *<* 1 (region III). However, at *σ* ≈ 3.4, there is a saddle–node bifurcation where endemic equilibria emerge and render the system bistable in the absence of control (above the red horizontal line in Fig. 6A). The coupled system remains bistable (with different alternative attractors), provided control costs are high. For sufficiently low costs, however, there will be successful disease eradication. In region I, this is because the control equilibrium disappeared in a transcritical bifurcation (leading to a phase portrait as in Fig. 4E). In region II, this is because the limit cycle disappeared in a homoclinic bifurcation (leading to a phase portrait as in Fig. 4D). The effect of infectivity enhancement is that disease eradication becomes more likely with increasing *σ*, as the white areas expand. Note also that increasing *σ* reduces the parameter domain where the endemic equilibrium is an attractor.

**Fig. 6.**
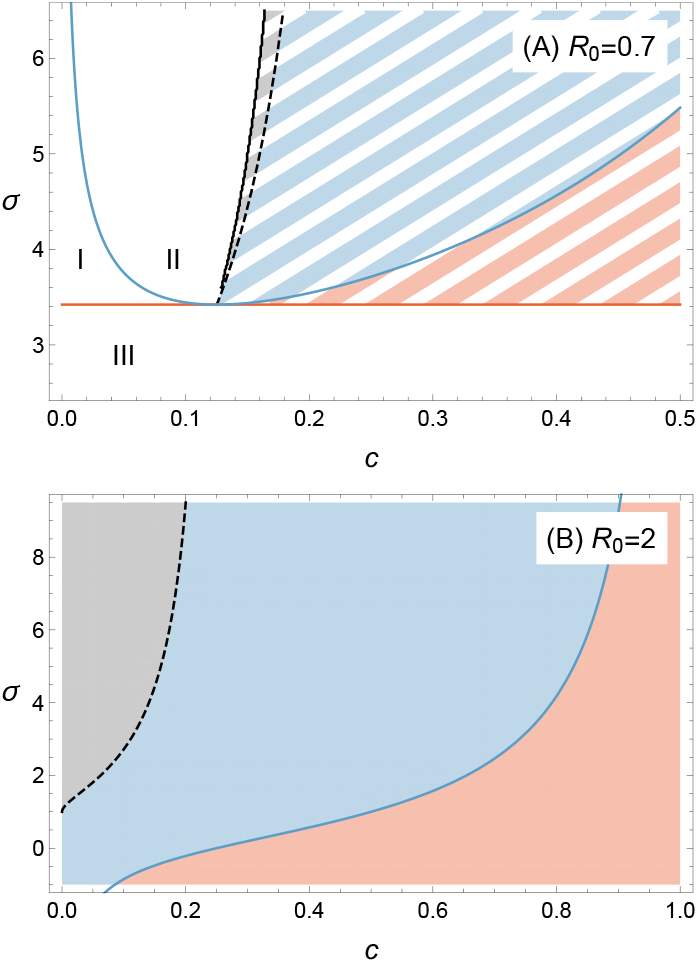
Dynamical regimes in the coupled model (14) in a two-parameter plot. White areas mark disease eradication, with the DFE being the only attractor. See the main text for the difference between regions I, II, and III. Hatched areas mark bistability. One of the attractors is always the DFE; the other attractor can be either a limit cycle (gray), the control equilibrium (blue), or an endemic equilibrium (red). In the monochromatic areas of (B), there is monostability with these attractors. The red line marks a saddle–node bifurcation curve where the two endemic equilibria coalesce and is given by (16). The blue curve marks transcritical bifurcations in which the control equilibrium (dis-)appears; it is given by (B.12). The dashed black curve marks Hopf bifurcations, which satisfy (B.17). The solid black curve marks homoclinic bifurcations and has been approximated numerically. Other parameter value: *b* = 1.

### 4.2 Monostable epidemiological scenario

Figure 6B shows a two-parameter bifurcation diagram when *R*_0_ *>* 1 and there is a unique stable endemic equilibrium in the absence of control. Increased infectivity enhancement promotes the uptake of control. It also promotes the destabilization of the system at low control costs. Hopf bifurcations can only occur for *σ >* 1 and 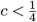, see Appendix B.3. If there is no Hopf bifurcation, control uptake monotonically increases with decreasing *c* (see the bifurcation diagram in Fig. 7A, *σ <* 1). If there is a Hopf bifurcation, the (equilibrium or time-averaged) control uptake shows a unimodal relationship with *c* (Fig. 7B, *σ >* 1); for details see Appendix B.3.

**Fig. 7.**
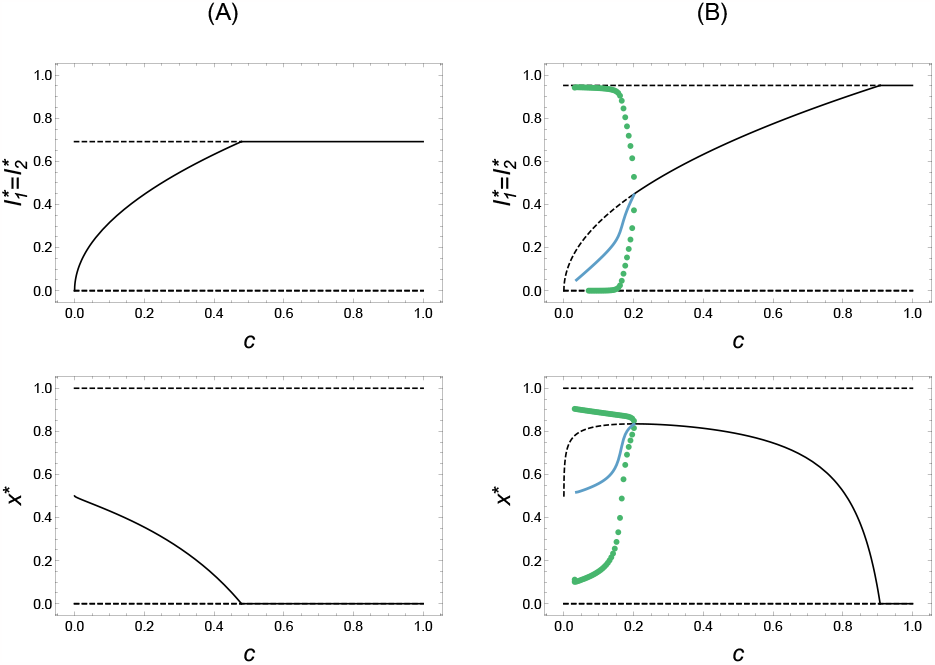
Bifurcation diagrams of the coupled model (14) when *R*_0_ *>* 1. Parameter values: *R*_0_ = 2, *b* = 1, (A) *σ* = 0.9 and (B) *σ* = 10. The meaning of lines and symbols is as in Fig. 5.

Last, consider the scaled learning rate *b*. The mathematical analysis in Appendix 4 reveals that it affects neither the existence nor the location nor the stability of any equilibrium point.

## 5. Discussion and conclusions

Game-theoretical and economic approaches to epidemiology generally predict that control coverage driven by self-interest is generally below the group-optimal level required for disease eradication (Fine & Clarkson, 1986; Bauch et al., 2003; Galvani et al., 2007; Fu et al., 2011; Chang et al., 2020). As control measures reduce disease prevalence they also incentivize individuals to become free-riders. Here, we have shown that selfish grower behavior can actually result in the socially optimal solution, namely the eradication of the disease.

There are only a few modeling studies offering solutions to the free-rider problem. Brito et al. (1991) analyzed conditions, in the form of subsidies or taxes, under which voluntary vaccination may be the better outcome than compulsory vaccination. Liu et al. (2012) found that the group-optimal control coverage can be reached by selfish individuals for diseases that become more severe with age, such as chickenpox. Investigating the effect of imperfect vaccines, Wu et al. (2011) found that selfinterested behavior can result in full coverage for intermediate vaccine efficacies. However, while the disease prevalence is reduced, the disease will not be eradicated. Our finding is, therefore, a rare exception that resolves the apparent impossibility of disease eradication under voluntary control programs.

The mechanism that enables disease eradication by self-interested behavior is the following. For diseases caused by coinfection, the disease may persist because of a positive feedback process (the more coinfections, the more transmission because of enhanced infectivity). Disease control undermines this feedback cycle that critically depends on a sufficiently large prevalence of coinfections. Essentially, control reduces the number of coinfections and thus counteracts the reinforcing cycle of enhanced infections. In some other sense, the disease control (if sufficiently affordable) transforms the basins of attraction such that the disease will be eradicated for all initial conditions. Mathematically, this can be reflected by a transcritical bifurcation, where the control equilibrium disappears (Fig. 6, region I), or by a homoclinic bifurcation, where the limit cycle oscillations push the system below the minimum disease prevalence required for coinfection-enhanced transmission (Fig. 6, region II).

There are ecological analogies of this mechanism. In populations subject to a strong Allee effect, cooperation between individuals can facilitate population persistence provided the population size is large enough (Courchamp et al., 2008). Harvesting this population can push the population below its minimum viable size, leading to population extinction (Clark, 1976; Segura et al., 2017; Hilker & Liz, 2020). Similar situations occur when a prey population is subject to both a strong Allee effect and predation (Bazykin, 1998; van Voorn et al., 2007) or a host population is subject to both a strong Allee effect and parasitism (Hilker et al., 2009; Hilker, 2010). Just like the strong Allee effect and harvesting, predation, or parasitism interact to form an ‘extinction vortex’, the coinfection-enhanced infectivity and disease control can jointly result in disease eradication.

The voluntary uptake of control needed for disease eradication is not very high. In fact, its asymptotic value will be nil. The situation is, therefore, essentially different from the conventional epidemiological scenarios where control uptake needs to exceed a herd immunity threshold that continually increases with *R*_0_. The requirement for disease eradication driven by self-interested behavior, however, is that control costs are sufficiently affordable (small *c*) and that the disease is sustained by coinfections (sufficiently large *σ* and *R*_0_ *<* 1).

There are many other epidemiological models that show bistability and disease persistence if *R*_0_ *<* 1. Examples include disease-induced mortality in vectorborne diseases, exogenous reinfection or imperfect vaccine in tuberculosis, differential susceptibility in risk-structured models, or vaccine-derived immunity waning at a slower rate than natural immunity. See Gumel (2012) and references therein, and also Hamelin et al. (2017, 2023) for more recent examples. All these models exhibit backward bifurcations and bistability, and it may be speculated whether self-interested behavior could also lead to disease eradication if these models were coupled with behavioral dynamics. An interesting aspect is that disease eradication in these epidemiological settings is often considered particularly difficult because protective measures need to reduce *R*_0_ significantly below one.

Our model is based on a number of simplifying assumptions. One is that the viruses are epidemiologically interchangeable, meaning that both pathogens have the same *R*_0_. Our model also implicitly assumes that intra-pathogen competition is greater than inter-pathogen competition. This means that pathogen 2 can always invade a population infected by pathogen 1 and the other way around (unless *σ* = −1, which is a limit case). Therefore, there is no priority effect, which would express itself as bistability between boundary equilibria (e.g. Gao et al., 2016). Another assumption is that only infectivity (rather than also susceptibility) is modified by coinfection. While this may be biologically plausible, it limits the generality of our model. Furthermore, our model assumed that hosts recover from the viruses, which, at least in plants, happens rarely. Last, but not least, we did not explicitly model the within- and between-field scales. Further work will be required to investigate how the results hold or change when more refined assumptions are taken into account.

On the other hand, these assumptions allowed us to formulate a very simple model. In fact, our coinfection model is the simplest model with multiple infections exhibiting bistability that we know (Keeling & Rohani, 2007; Gao et al., 2016; Allen et al., 2019; Chapwanya et al., 2021). The usefulness of our approach is that we obtained a very simple (two-dimensional) coinfection model, into which we could integrate behavioral dynamics. The coupled model has only four parameters. This allowed us to obtain a fairly comprehensive understanding of the system dynamics.

It turned out that the scaled learning rate *b* has no effect on existence, location, and stability of equilibria. However, it does have an effect on the speed with which attractors are approached, and on the limit cycle oscillations. Parameter *b* essentially describes the relative time scales between the behavioral and the epidemiological dynamics. In contrast to our results, Bauch (2005) found that the scaled learning rate in his pertussis model did affect the Hopf bifurcation curve, for example. In social-ecological models of shallow lake eutrophication, where the behavioral dynamics are described by stochastic best-response equation, there is a similar contrast. In some models, a similar time scale parameter was found to affect the stability (Suzuki & Iwasa, 2009; Iwasa et al., 2010), whereas it was found to have no effect, as in the current study, in Sun & Hilker (2020).

The occurrence of limit cycles (or “wave-like oscillations”) is well known from behavioral epidemic models (Bauch, 2005; d’Onofrio et al., 2007; Poletti et al., 2009; Bhattacharyya & Bauch, 2010; Zhang et al., 2010; Cornforth et al., 2011). What is notable here is that the time-averaged control coverage decreases as control becomes less expensive. This may seem counter-intuitive. However, it probably reflects the fact that the cycle is slowed down when control and disease prevalence are low. Moreover, there is an analogy to the “hydra effect” in ecology, where increased predator mortality can increase mean predator population size in predator–prey cycles (Abrams, 2009; Sieber & Hilker, 2012).

Overall, we have shown that the optimal outcome at the group level (i.e., disease eradication) can be achieved by “rational” individuals (i.e., growers voluntarily adopt control measures to maximize their payoff) – even in the absence of institutional design principles (Ostrom, 1990) or cooperative mechanisms (e.g., kin or group selection or a form of reciprocity, Nowak, 2006). This occurs in a bistable epidemiological context with reinforcing processes, and when control costs are sufficiently low. Alternative stable states, regime shifts, and tipping points are prevalent in many environmental (Scheffer et al., 2001), climate (Lenton et al., 2008), and social systems (Nyborg et al., 2016). It therefore remains an intriguing question whether similar resolutions of social or public good dilemma might exist in such systems as well and offer pathways to more desirable system states.

## Acknowledgments

We acknowledge partial funding from the ANR project BEEP (Behavioral Epidemiology and Evolution of Plant Pathogens).

## Appendix A. Analysis of the coinfection model

The coinfection model, i.e., Eq. (17), is repeated here for convenience:

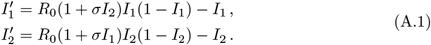

The biological domain of (*I*_1_, *I*_2_) is [0, 1]^2^.

### A.1 Equilibria and their existence in the biological domain

The model has at most five equilibria:

- the disease-free equilibrium, (0, 0), which always exists;
- two symmetric mono-pathogenic equilibria, (1 − 1*/R*_0_, 0) and (0, 1 − 1*/R*_0_), which exist if and only if *R*_0_ *>* 1;
- two endemic equilibria in which both pathogens coexist,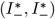 and 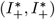, which we study in more detail below.

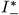 and 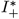 are the possible solutions of the following quadratic equation:

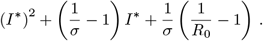

Thus,

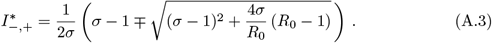

The term under the square root is positive if and only if

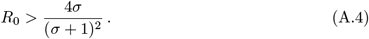

If *σ <* 0, this condition is always satisfied. Note also that since

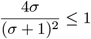

(with equality for *σ* = 1), condition (A.4) is always satisfied when *R*_0_ *>* 1. Condition (A.4) may not be satisfied only when *σ >* 0 and *R*_0_ *<* 1.

Next, one can show that

1. 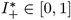 if and only if *σ* > 1, or 0 < *σ* < 1 and R_0_ > 1,
2. 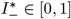 if and only if *σ* > 1 and R_0_ < 1, or −1 < *σ* < 0 and R_0_ > 1 Altogether, the existence results are summarized in Tab. 2. Note that the endemic equilibria are ordered in the sense that 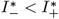and that this applies to the infection compartments with both virus 1 and 2. We shall refer to these equilibria as the larger and smaller endemic equilibria.

### A.2 Stability analysis

The Jacobian of model (A.1) is:

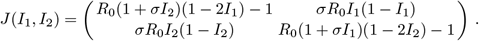

#### A.2.1 Disease-free equilibrium

The Jacobian evaluated at the disease-free equilibrium is:

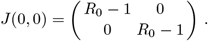

It has the two identical eigenvalues *λ*_1_ = *λ*_2_ = *R*_0_ − 1. Therefore, the disease-free equilibrium is locally asymptotically stable if and only if *R*_0_ *<* 1.

#### A.2.2 Mono-pathogenic equilibria

We evaluate the Jacobian at (1 − 1*/R*_0_, 0), which is one of the two symmetric monopathogenic equilibria:

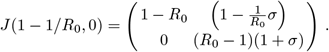

Since det(*J*) = −(*R*_0_ − 1)^2^(1 + *σ*) *<* 0, this mono-pathogenic equilibrium is an unstable saddle point. The same holds for the other mono-pathogenic equilibrium due to the symmetry.

#### A.2.3 Endemic equilibria

We first notice that the monotone dynamical system (A.1) is cooperative (competitive) if and only if *σ >* 0 (*σ <* 0, respectively) (Hirsch & Smith, 2006). Since the system is two-dimensional, and the square [0, 1]^2^ is invariant, the solution converges to an equilibrium, i.e., there is no stable periodic orbit (Hirsch, 1982).

If *R*_0_ *>* 1, the disease-free equilibrium is unstable and the solution thus converges to the endemic equilibrium 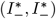 (if *σ <* 0), or 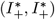 (if *σ >* 0) (see Tab. 2), which is therefore globally asymptotically stable.

If *R*_0_ *<* 1 and *σ <* 1, or *σ >* 1 and *R*_0_ *<* 4*σ/*(*σ* + 1)^2^, there is no endemic equilibrium (Tab. 2), and the solution converges to the disease-free equilibrium, which is therefore globally asymptotically stable.

The remaining case is *σ >* 1 and 4*σ/*(*σ* + 1)^2^ *< R*_0_ *<* 1, in which two endemic equilibria, 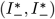 and 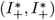, exist in the biological domain (Tab. 2). We have

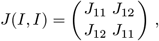

in which

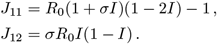

The eigenvalues of *J* are such that

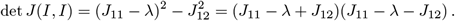

Therefore,

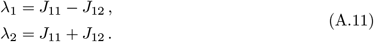

Since *J*_12_ *>* 0, *λ*_1_ *< λ*_2_.

We now treat the larger and smaller endemic equilibrium separately:

1. *Larger endemic equilibrium*. Using 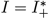 as given by Eq. (A.3), we obtain:

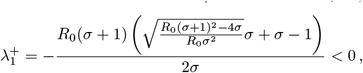

since we assume *σ >* 1 and *R*_0_ *>* 4*σ/*(*σ* + 1)^2^ in this case. We also obtain

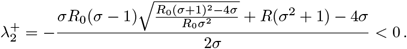

This means that the larger endemic equilibrium, 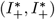, is locally asymptotically stable when it exists.
2. *Smaller endemic equilibrium*. Using 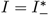 as given by Eq. (A.3), and using the fact that *R*_0_ *<* 1, one can easily show that:

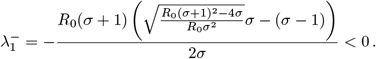

Similarly,

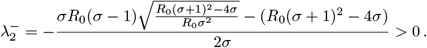

This means that the smaller endemic equilibrium 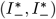 is an unstable saddle point when it exists. Since *R*_0_ *<* 1, the disease-free equilibrium is locally asymptotically stable. Therefore, there is bistability between the disease-free equilibrium, (0, 0), and the larger endemic equilibrium, 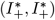. More specifically, the dynamics converge to one or the other equilibrium depending on the initial conditions (see Fig. 8).

**Fig. 8.**
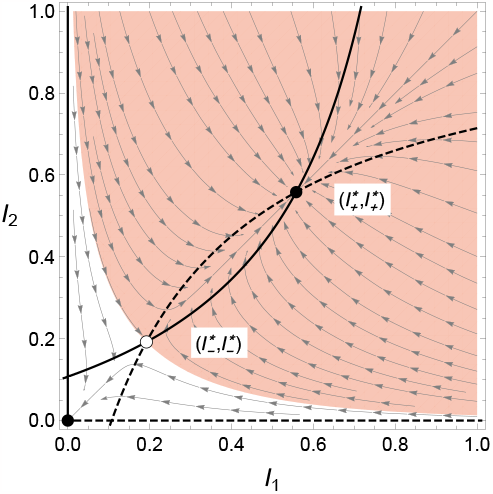
Phase plane portrait of the coinfection model (15). Stable (unstable) equilibria are shown as filled (empty, respectively) circles. The solid (dashed) curves are the nullclines of *I*_1_ (*I*_2_, respectively). The white (red) areas are the basins of attraction to the disease-free equilibrium (larger endemic equilibrium, respectively). Parameter values: *R*_0_ = 0.7, *σ* = 4.

## Appendix B. Analysis of the coupled model

In the following, we will find that there can exist up to seven equilibria 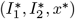 of the coupled system (14). We will investigate their local stability with the help of the Jacobian, which reads

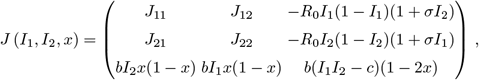

Where

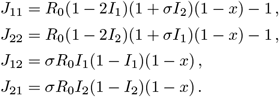

### B.1 Equilibria with simple conditions for existence and stability

Two equilibria exist unconditionally. One is *E*_0_ = (0, 0, 0), to which we shall refer as the disease-free equilibrium (DFE). The eigenvalues of the Jacobian evaluated at the DFE are

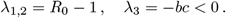

The latter conveys that the DFE is locally attractive in the *x*-dimension of the phase space. Overall, the DFE is a stable node if and only if *R*_0_ *<* 1 and an unstable saddle point otherwise.

The other unconditionally existent equilibrium is *E*_F_ = (0, 0, 1), to which we shall refer as the full-control equilibrium. The eigenvalues are

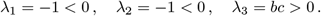

The full-control equilibrium is always an unstable saddle point.

There are two equilibria that conditionally exist depending on only *R*_0_. They are the mono-pathogenic equilibria 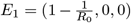 and 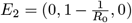 with virus 1 and 2, respectively. They both exist if and only if *R*_0_ *>* 1. For *E*_1_, the eigenvalues are

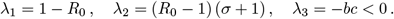

If *E*_1_ exists, i.e., *R*_0_ *>* 1, then *λ*_2_ *>* 0 and *λ*_1,3_ *<* 0. Therefore, it is always an unstable saddle point.

For *E*_2_, the eigenvalues are

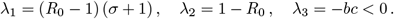

If *E*_2_ exists, i.e., *R*_0_ *>* 1, then *λ*_1_ *>* 0 and *λ*_2,3_ *<* 0. Therefore, it is always an unstable saddle point.

### B.2 Endemic equilibria

There are two equilibria that conditionally exist depending on only the epidemiological parameters *R*_0_ and *σ*. They are

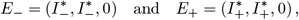

where 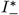 and 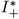 are as in Eq. (A.3) of the coinfection model. The existence conditions are the same as the ones summarized in Tab. 2. We shall refer to *E*_−_ and *E*_+_ also as the smaller and larger endemic equilibrium, respectively, but note that they are pure equilibria in a game-theoretic sense because all growers adopt the same strategy of no control.

Now let us turn to the stability analysis of the endemic equilibria in the coupled model. First of all, we acknowledge that *E*_−_ and *E*_+_ lie in the plane *x* = 0 and on the diagonal given by *I*_1_ = *I*_2_. We will focus on this property and evaluate the Jacobian at points *P* = *{*(*I*_1_, *I*_2_, *x*) |*x* = 0, *I*_1_ = *I*_2_*}*. Referring to *I*_1_ and *I*_2_ as *I* due to their equal values at any of the two endemic equilibria, the Jacobian for points *P* is:

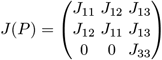

With

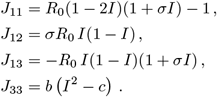

In order to determine the nontrivial eigenvalues, the characteristic polynomial needs to be zero:

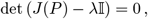

where 𝕀 denotes the identity matrix. Developing the determinant, we obtain

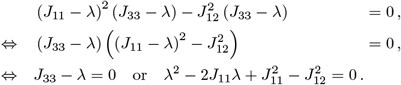

The eigenvalues of the Jacobian at points *P* are thus:

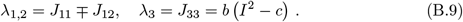

*λ*_1_ and *λ*_2_ are the same as for the endemic equilibrium in the pure coinfection model, cf. Eq. (A.11). Therefore, the smaller endemic equilibrium in the coupled model, *E*_−_, has 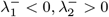 and is always an unstable saddle point, independent of *λ*_3_. The larger endemic equilibrium in the coupled model, *E*_+_, has 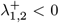. Hence, it is locally asymptotically stable if and only if *λ*_3_ *<* 0, which is equivalent to

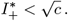

Since 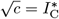, this means that the larger endemic equilibrium is locally asymptotically stable if and only if its infection levels are smaller than those of the control equilibrium.

Note that parameter *b* does not influence the local stability of the endemic equilibria. This is because *b* is not included in *λ*_1,2_ and cannot change the sign of *λ*_3_.

### B.3 Control equilibrium

Finally, there can be a unique nontrivial equilibrium 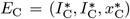 with the components

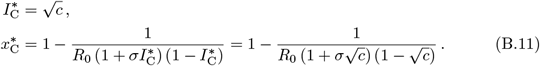

We shall refer to it as the control equilibrium. Note that it is the only “mixed” equilibrium in a game-theoretic sense.

Its existence requires, on the one hand, 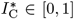, which is equivalent to *c* ∈ [0, 1]. On the other hand, its existence requires 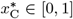. First, consider 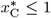, from which we obtain

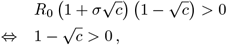

which is always true when 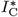exists, i.e., for *c* ∈ [0, 1]. Second, 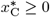 if and only if

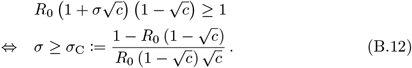

Condition (B.12), together with *c* ∈ [0, 1], guarantees the existence of the control equilibrium. Figure 9 shows how *σ*_C_ depends on *c* and *R*_0_. If *R*_0_ *>* 1, there is a unique value of *c*, below which the control equilibrium exists. If *R*_0_ *<* 1, and for sufficiently large infectivity enhancement *σ*, there are two values of *c*, in between the control equilibrium exists, i.e., the control equilibrium disappears for small enough *c*.

**Fig. 9.**
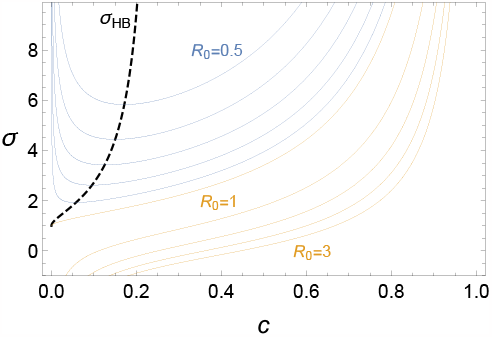
Existence and stability conditions for the control equilibrium. The control equilibrium exists in the parameter regions above the solid curves, which are shown for different values of *R*_0_ (blue: *R*_0_ between 0.5 and 0.9 in steps of 0.1; orange: *R*_0_ between 1 and 3 in steps of 0.5). The solid curves are *σ*_C_, as given by (B.12). The dashed black line marks the Hopf bifurcation curve and is given by *σ*_HB_ in (B.17), provided the control equilibrium exists. Limit cycles are possible in the parameter domain above the curve. Numerical simulations show that the limit cycles can disappear in homoclinic bifurcations (e.g., see Figs. 5 and 6).

Let us now investigate how parameter *c* influences the location of the control equilibrium. Clearly, 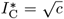 increases with *c* in form of a parabolic branch. The control coverage, 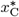 can be strictly monotonically decreasing with *c* (see, e.g., Fig. 7A) or can be a hump-shaped function of *c* (see, e.g., Fig. 7B). The latter occurs when 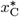 has a local maximum. Solving the extremum condition 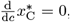, we obtain

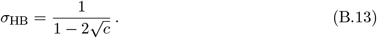

We will later see that this corresponds to a Hopf bifurcation condition.

Let us now investigate the local stability of the control equilibrium. The Jacobian reads

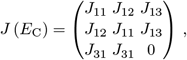

Where

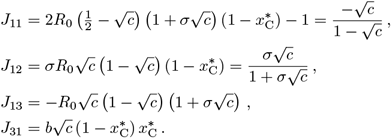

With the help of a computer algebra system, we find one of the three eigenvalues:

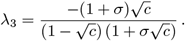

One can show that *λ*_3_ *<* 0 for all *c* ∈ (0, 1) and *σ >* −1. Hence, the stability of the control equilibrium depends only on *λ*_1,2_. They are

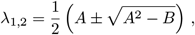

Where

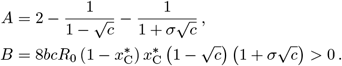

When substituting 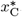, it is possible to find also explicit (but cumbersome) expressions for *λ*_1,2_. However, we are mostly interested in the occurrence of a Hopf bifurcation. We expect it to take place when *λ*_1,2_ are complex conjugate and their real parts change signs. From the latter, we can find 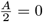 as a necessary condition for a Hopf bifurcation of the control equilibrium, which gives

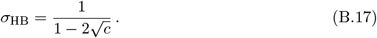

This coincides with the expression in Eq. (B.13), i.e., if there is a Hopf bifurcation, it takes place at the local maximum of 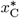 as a function of *c* (for example, see the lower panel of Fig. 7B).

Interestingly, condition (B.17) depends only on *c* and is independent of *R*_0_ and *b*. Figure 9 shows the graph. A prerequisite for the Hopf bifurcation to occur is *σ*_HB_ *>* −1, from which we obtain

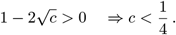

Furthermore, when 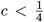, then *σ*_HB_ *>* 1. Taken together, Hopf bifurcations are only possible in the parameter domain

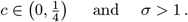

Finally, we find that parameter *b* does not affect the stability of the control equilibrium. *b* shows up only in the discriminants of *λ*_1,2_. If the discriminants are negative, the real parts of *λ*_1,2_ are 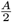 and thus independent of *b*. Alternatively, if the discriminants are non-negative, then *λ*_1,2_ are real and have the same signs, which are also independent of *b*.

